# Infection of the bovine mammary gland by avian H5N1 subclade 2.3.4.4b influenza viruses

**DOI:** 10.64898/2026.04.16.718897

**Authors:** Rebecca A. Ross, Sarah K. Walsh, Hannah Montgomery, Hanting Chen, Edward Hutchinson, Pablo R. Murcia

## Abstract

The emergence of the panzootic clade of highly pathogenic avian influenza H5N1 (2.3.4.4b) in 2020 marked a major expansion in the host range of influenza A viruses (IAVs), raising concerns about further cross-species transmission events and zoonotic spillover. Introduction of 2.3.4.4b viruses into U.S. dairy herds has resulted in widespread circulation, accompanied by reduced milk yield, mastitis, and high viral loads in milk. Notably, virus circulation in dairy cattle represents a novel route for mammalian adaptation and transmission that has already led to more than 40 human cases in the U.S. since 2024. Here, we investigated whether avian clade 2.3.4.4b viruses could infect mammary tissue from Aberdeen Angus, Holstein Friesian, and Limousin cattle, three breeds commonly farmed in Europe, the Americas, and Oceania. Using mammary gland explants, we inoculated tissues with attenuated reassortant viruses expressing the haemagglutinin and neuraminidase glycoproteins of three 2.3.4.4b viruses that predated the emergence of H5N1 in US cattle: A/chicken/England/053052/2021 (AIV07), A/chicken/Scotland/054477/2021 (AIV09), and A/chicken/England/085598/2022 (AIV48). Infected epithelial cells were identified using immunohistochemistry in explants from both the teat and gland cistern for all three breeds following infection with AIV09 and AIV48, indicating that mammary tissue from each of the three tested cattle breeds cattle is permissive to H5N1 infection. Lectin staining showed expression of both α2,3-linked and α2,6-linked sialic acids in the mammary tissue of all donors showing that all three breeds have the potential to support infection with both avian-adapted and mammalian adapted IAVs. Together, these findings demonstrate that mammary glands from both beef and dairy cattle breeds are permissive to infection with avian-adapted and mammalian-adapted H5N1 viruses and highlight the potential for this tissue to act as a mixing vessel for IAV reassortment, underscoring the need to include cattle in ongoing H5N1 surveillance and risk-assessment frameworks.

**Impact Statement:** The emergence of highly pathogenic avian influenza H5N1 in dairy cattle has expanded the recognised host range of influenza A viruses. Further, the ability of the virus to infect the mammary gland and transmit via milk revealed a novel interface for transmission to humans and animals. Although sustained circulation in US dairy herds has been reported, the susceptibility of mammary tissue from other breeds (including beef cattle) commonly used in different countries has been largely unexplored. Here, we show that avian-origin H5N1 viruses can infect tissues derived from the mammary gland of three common cattle breeds (Aberdeen Angus, Holstein Friesian, and Limousin). Virus was detected in epithelial cells from both dairy and beef breeds, indicating that H5N1 can infect multiple breeds. Receptor profiling showed abundant α2,3-linked and α2,6-linked sialic acids, consistent with a tissue environment that may support infection with both avian-adapted and mammalian-adapted viruses. These findings demonstrate that multiple cattle breeds are permissive to H5N1 infection and strengthens the evidence base for including cattle in H5N1 surveillance and risk-assessment frameworks.

## Introduction

Goose/Guangdong-lineage (Gs/Gd) H5N1 highly pathogenic avian influenza (HPAI) viruses were first detected in 1996 in domestic geese in Guangdong, China, and quickly established themselves as a major cause of mortality in poultry and wild birds across Asia [1,2]. Over subsequent years long-distance migration and reassortment drove their global expansion, with detections across >80 countries by the mid-2010s [3]. A critical transition occurred with the rise of clade 2.3.4.4 viruses, which exhibit high genetic plasticity and frequent reassortment, yielding multiple H5Nx constellations [3,4]. Within this group, clade 2.3.4.4b became predominant from 2019 onwards, with global dissemination and expanded host range – characterised by extensive circulation in wild birds and repeated spillovers into a variety of mammals, including explosive outbreaks in pinnipeds and mustelids [5–12]. Most recently, detection of H5N1 infection in dairy cattle in the United States [13,14] and the Netherlands [15] has underscored the threat of H5N1 spillover for both agriculture and public health.

Many factors influence the ability of influenza A viruses (IAVs) to infect different host species [16]. Primarily, susceptibility to infection with IAVs is determined by the presence of host cell receptors to which the virus binds and enter cells. For IAV, receptor usage is predominantly α2,3-linked sialic acids in relevant avian tissues and α2,6-linked sialic acids in humans. Other mammals exhibit varying balances of these receptors across tissue types, and these differences – alongside other host- and virus-specific factors – contribute to variation in host and tissue susceptibility to IAV infection. In cattle, recent H5N1 infections appear to show a marked tissue tropism for the mammary gland [13,17–20]. Affected cattle typically show mammary infection, with associated decreases in milk production, mastitis, and high viral loads in milk [13,17,18]. By contrast, reports of respiratory infection or respiratory signs remain limited [13,18,19,21]. Observations of limited viral RNA in nasal swabs alongside high infectious titres in oral swabs and milk in affected herds support a model in which mammary infection and milk contamination play central roles in onward spread in dairy settings [22]. Sustained replication of H5N1 in the bovine mammary gland may therefore exert tissue-specific selection pressures that differ from those of the respiratory tract. The consequences of this selective environment for virus replication, transmission, and cross-species infection remain unknown.

Here, we evaluated the permissiveness of common beef and dairy cattle to different avian-origin 2.3.4.4b H5N1 isolates, to assess the risk of H5N1 infection in agricultural settings. We infected *ex vivo* cultures of mammary glands from six donors with attenuated reassortant viruses expressing the haemagglutinin (HA) and neuraminidase (NA) of the H5N1 viruses A/chicken/England/053052/2021 (AIV07), A/chicken/Scotland/054477/2021 (AIV09), and A/chicken/England/085598/2022 (AIV48) or the HA and NA of human seasonal viruses (A/Norway/3433/2018 H1N1 and A/Norway/3275/2018 H3N2) and identified infected cells by immunohistochemistry. We additionally characterised the distribution of α2,6- and α2,3-linked sialic acids in donor tissues using lectin staining.

## Methods

### Biosafety considerations

All experiments with attenuated reassortant viruses were approved by the local genetic manipulation safety committee at the University of Glasgow (GM223), and the Health and Safety Executive of the United Kingdom. All experiments were carried out in appropriate laboratories by trained personnel at biosafety level 2 (BSL2). To allow us to investigate virus tropism, all attenuated reassortant viruses used in this study were derived by reverse genetics using the internal genes (PB2, PB1, PA, M, NP, M, and NS) of the laboratory-attenuated vaccine strain A/Puerto Rico/8/34 (H1N1) and the glycoproteins (HA and NA) of the virus of interest. Additionally, all HA genes from highly pathogenic avian influenza viruses (HPAIVs) had the polybasic cleavage site (a major contributor to viral pathogenicity) removed. Work with the attenuated reassortant viruses used in this study was physically segregated from work all other work with replication-competent IAVs to prevent reassortment generating viruses with novel HA/NA combinations.

### Ethics statement

All animal work conducted was in accordance with the animal ethics and welfare committee at the University of Glasgow. Work using animal tissues was approved by the Ethics Committee of the School of Veterinary Medicine of the University of Glasgow (ethics approval EA26/25).

### Collection of bovine mammary

Udder tissue derived from six female cattle (Table S1) was collected from commercially slaughtered cows free from any antibiotic treatment or mastitis. The teat and gland cistern were removed from each udder quarter (Figure S1) and transferred into Dulbecco’s Modified Eagles Medium (DMEM, Gibco) supplemented with 2% penicillin-streptomycin (Gibco) and 2% amphotericin B (Gibco), hereafter referred to as mammary explant growth media. Using a 5mm biopsy punch (Covetrus), tissue cores (explants) were prepared from each tissue type and stored in mammary explant growth media until infection.

### *Ex vivo* culture

Mammary explants were prepared and maintained in culture as previously described [23]. Briefly, explants were placed in 96-well plates and submerged in mammary explant growth media supplemented with 10% heat-inactivated foetal bovine serum (FBS, Gibco). Plates were cultured overnight in a humidified incubator at 37°C with 5% CO2.

### Explant RNA extractions and pan-IAV RT-qPCR

All tissue RNA extractions were performed using a modified version of the RNeasy Plus RNA Extraction Kit (Qiagen). In brief, two tissue samples for each unique combination of donor and anatomical location were placed in separate 1.5 mL cryovials alongside 600 uL of Buffer RLT (Qiagen) containing beta-mercaptoethanol (ThermoFisher) and one 7 mm stainless steel bead (Qiagen). The samples were homogenised using a Precellys 24 Tissue Homogeniser (Bertin Technologies) set to 4000 rpm for 90 s a total of three times. The supernatant was then transferred to a gDNA Elimination Column (Qiagen) and the extraction proceeded as per the manufacturer’s instructions.

Following the RNA extraction, a pan-IAV RT-qPCR was performed using a QuantStudio 3 Real-Time PCR (qPCR) system (Applied Biosystems) to ensure that the samples were not infected with IAV prior to the experimental infection. In brief, the v2 Nagy M-gene primers [24] were used alongside the QuantiNova Probe RT-PCR kit (Qiagen) with all conditions as per the manufacturer’s instructions except for the annealing temperature, which was changed to 58°C. Two explants from each tissue and donor were tested with two technical replicates of each sample run per RT-qPCR plate. All plates were run with three NTCs and three linearised M-gene positive controls. Samples were considered negative if no amplification curve was observed within 40 cycles. All tested samples were observed to be negative.

### Cells and viruses

Wild-type A/Puerto Rico/8/34 (PR8) was generated in HEK293T cells using the pDUAL reverse genetics system, as previously described [25]. HA and NA plasmids for both the H5N1 and seasonal viruses used here were a kind gift from Professor Munir Iqbal (The Pirbright Institute). Attenuated reassortant viruses with the glycoproteins of either a H5N1 virus or seasonal IAV and the internal genes of PR8 were also generated in HEK293T cells using the pDUAL reverse genetics system (Table 1). Viruses were passaged at a low multiplicity of infection (MOI) in MDCK cells in viral growth media (VGM) consisting of DMEM (Gibco) with 1 μg/μL TPCK-treated trypsin (Thermo) until approximately 70-80% CPE was observed. At this point the supernatant was clarified by centrifugation and stored at −80°C in small volume aliquots. Frozen aliquots were only used in experimental infections to avoid freeze-thaw changes to infectious titre. Viral titre was determined using two methods: 1) plaque assays to determine plaque forming units per millilitre (PFU/mL) [26]; and 2) RT-qPCR to determine genome copies per millilitre (copies/mL) (see RT-qPCR section).

**Table 1:** Viruses used in this study. For all strains indicated, reassortant viruses were produced with the HA and NA genes of the indicated viruses and the remaining genes from the PR8 strain. For AIV strains the polybasic cleavage site of HA was replaced by a monobasic cleavage site.

| Short name | Isolate ID | Subtype |
| --- | --- | --- |
| PR8 | A/Puerto Rico/8/34 | H1N1 |
| H1N1 | A/Norway/3433/2018 | H1N1 |
| H3N2 | A/Norway/3275/2018 | H3N2 |
| AIV07 | A/chicken/England/053052/2021 | H5N1 |
| AIV09 | A/chicken/Scotland/054477/2021 | H5N1 |
| AIV48 | A/chicken/England/085598/2022 | H5N1 |

### Phylogenetic inference

To show the evolutionary relationships between the H5N1 isolates used in this study and other H5N1 isolates, a maximum likelihood (ML) phylogeny was inferred. Additional isolates included: A/goose/Guangdong/1/1996 (EPI_ISL_2584399), A/dairy cow/Texas/24-008749-001-original/2024 (EPI_ISL_19014384), Human A/Texas/37/2024 (EPI_ISL_19027114), A/dairy cow/USA/002645-005/2025 (EPI_ISL_19716907), and A/British_Columbia/PHL-2032/2024 (EPI_ISL_19548836). The phylogeny was inferred using PhyML v3.3.20180621 implemented within Geneious. General Time Reversible (GTR) substitution model with a gamma-distributed rate variation (+G, 4 categories) and a proportion of invariant sites (+I) was applied. Model parameters (transition/transversion ratio, gamma shape parameter, and proportion of invariant sites) were estimated from the data, and tree topology, branch lengths, and substitution rates were jointly optimized. Branch support was assessed using 1,000 non-parametric bootstrap replicates. The resulting unrooted tree was subsequently rooted using the A/Norway/3275/2018 H3N2 HA sequence as an outgroup.

### Experimental infections

Media were removed from the mammary explants prior to infection. Viruses were diluted to 1×10^9^ genome copies per mL (measured using the pan-IAV RT-qPCR assay) and 5 uL was used to infect the apical surface of each explant for a final dose of 5×10^6^ genome copies per infection. Two explants were infected per donor for each virus. After 1 hour of incubation, the explants were re-immersed in mammary growth media. Plates were cultured for 24 hours in a humidified incubator at 37°C with 5% CO2. Explants were then fixed in 10% formaldehyde at 4°C for 24 hours, dehydrated in 70% ethanol, and embedded in paraffin wax to generate formalin-fixed and paraffin-embedded (FFPE) tissue blocks. 2–3 μm longitudinal sections were taken for immunohistochemistry (IHC) and to assess epithelial integrity by haematoxylin and eosin (H&E) staining. For H&E staining, sections were deparaffinized and rehydrated prior to staining.

### Immunohistochemistry

FFPE tissue blocks were cut into 2–3 μm sections as described above and mounted on glass slides, before undergoing deparaffinization and rehydration [27]. For immunohistochemistry (IHC), sections were antigen-retrieved in pH 6.0 citrate buffer solution at 90°C for 30 min. Sections were washed three times in 1X Phosphate-Buffered Saline solution (PBS, Thermo Fisher). To block endogenous peroxide activity, slides were incubated in PBS with 3% H202 solution (Thermo Fisher) at 4°C for 20 min, washed three times with PBS, and blocked in 1% bovine serum albumin (BSA, Thermo Fisher) for 30 minutes at room temperature. Sections were then incubated with DA183 sheep anti-IAV-NP antibody (European Veterinary Laboratory, Clone HB65) diluted 1:500 in PBS with 1% BSA overnight at 4°C. Finally, sections were washed again before being incubated in Biotin-SP-conjugated AffiniPure donkey anti-sheep IgG (Jackon ImmunoResearch) diluted 1:500 in PBS with 1% BSA for 1 hour at room temperature (RT). Sections were incubated with extravidine peroxidase (Sigma-Aldrich) diluted 1:100 in PBS with 1% BSA for 1 hour at RT. 3,3’-Diaminobenzidine (DAB, Merk) was used as a chromogen and Mayer’s hemalum solution (Sigma Aldrich) was used as a counterstain in all IHC experiments. Slides were subsequently dehydrated and mounted as previously described [27].

### Lectin staining

Tissue sections were prepared as described above. Antigen retrieval was performed by incubating sections in 0.01 M EDTA for 30 min at 95°C before cooling to RT. Sections were then blocked for 30 min in PBS with 1% BSA and washed three times with PBS before incubating overnight at 4°C with 20 μg/mL of either Fluorescein-conjugated MAL-I or CY5-conjugated SNA (both Vector Laboratories) diluted in PBS with 1% BSA. Sections were treated with streptavidin conjugated with Alexa Fluor 488 (Invitrogen) and incubated for 1 hour at 4°C. Finally, slides were stained with Hoechst (Thermo Fisher) diluted 1:500 for 30 minutes and mounted with ProLong™ Gold Antifade (Invitrogen). Images were captured using a Zeiss LSM 710 microscope with Fast Airyscan at 63x magnification.

### Digital pathology

For IHC, slides were digitalised and scanned on the Zeiss Axioscan 7 using the Plan-Apochromat 40x/0.8 objective. Image processing was conducted using Zeiss ZEN software (v3.10), with uniform exposure, gain, brightness, and contrast for each tissue type.

## Results

### Bovine mammary tissue displays α2,3-linked and α2,6-linked sialic acids

To assess the display of IAV receptors in bovine mammary tissue, lectin staining was performed on FFPE blocks containing teat and gland cistern explants from Aberdeen Angus, Holstein Friesian, and Limousin cattle. Staining with the lectins MAL-1 and SNA revealed widespread display of both α2,3-linked and α2,6-linked sialic acids in the epithelial cells of both the gland and teat cisterns across all three breeds (Figure 1). This signal was predominantly localised to the luminal surface of epithelial cells, with the labelling presenting as multifocal to diffuse. No qualitative differences in staining distribution or intensity were observed between beef and dairy breeds, or between lactating or non-lactating donors, however there was significant between-donor variation both within and across breeds (Table 2). These data demonstrate that bovine mammary tissue displays both α2,3-linked and α2,6-linked sialic acids, with broadly similar distributions across breeds and physiological states.

**Table 2:** Lectin staining results for all donors. The table summarises the localisation of signals representing lectin staining results for each of the 6 donors used in this study. Samples are only reported as not showing expression if all biological and technical replicates of that unique combination of donor and tissue were negative. Cattle breeds are indicated with ‘AA’ for Aberdeen Angus, ‘L’ for Limousin, and ‘HF’ for Holstein Friesian, individual donors are identified as D1-D6. Staining intensity was expressed by relative grades from negative (-) to weak (+) to strong (++), see Figure S2 for representative images of each relative grade. MAL 1 = α2,3, SNA = α2,6.

|  |  |  | Apical surface |  | Epithelial layer |  |
| --- | --- | --- | --- | --- | --- | --- |
| | | | α2,3 | α2,6 | α2,3 | α2,6 |
| AA | D1 | Teat | + | ++ | + | ++ |
|  |  | Gland | + | ++ | + | ++ |
|  | D2 | Teat | + | ++ | ++ | ++ |
|  |  | Gland | + | ++ | ++ | ++ |
|  | D6 | Teat | + | ++ | + | ++ |
|  |  | Gland | + | ++ | + | + |
| L | D3 | Teat | ++ | ++ | ++ | + |
|  |  | Gland | - | ++ | + | + |
|  | D4 | Teat | ++ | ++ | ++ | - |
|  |  | Gland | ++ | ++ | ++ | + |
| HF | D5 | Teat | + | ++ | + | ++ |
|  |  | Gland | ++ | ++ | ++ | ++ |

**Figure 1:**
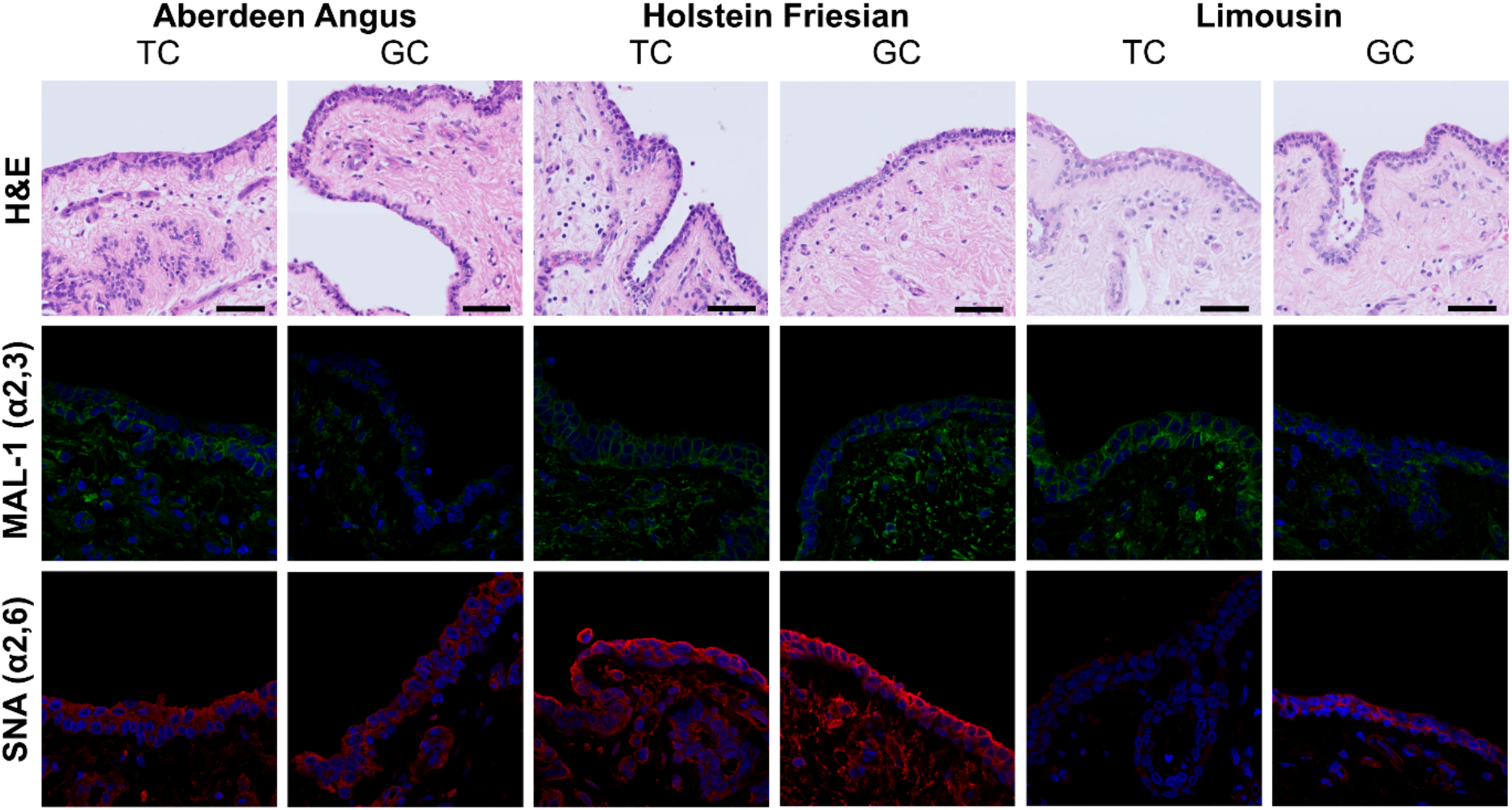
Lectin staining results for one representative donor of each cattle breed. The panels show Haematoxylin and Eosin (H&E) staining, MAL-1 staining for α2,3-linked sialic acids, and SNA staining for α2,6-linked sialic acids for one donor of each breed for both the teat cistern (TC) and gland cistern (GC). For the lectin staining, α2,3-linked sialic acids are shown in green, α2,6-linked sialic acids in red, and cell nuclei are shown in blue (DAPI stained). Donor 2 is shown for Aberdeen Angus, Donor 5 for Holstein Friesian, and Donor 4 for Limousin.

### Bovine mammary tissue is permissive to H5N1 clade 2.3.4.4b infection

To determine whether bovine mammary tissue is permissive to IAV infection, explants were inoculated with attenuated reassortant viruses bearing the HA and NA of three clade 2.3.4.4b H5N1 isolates (AIV07, AIV09, AIV48; phylogenetic context shown in Figure 2), alongside human seasonal H1N1 and H3N2, PR8, and mock controls. After 24 hours the explants were fixed and viral antigen detected by immunohistochemistry. Viral antigen was detected in multiple explants across breeds, although the extent of infection varied by virus, donor, and tissue type (Figure 3 and Table 3). Generally, donors were more permissive to infection in the teat cistern, with only limited infection observed in the gland cistern (D1-D4). Similarly, donors were more permissive to infection with some viruses than others, with all donors showing infection in at least one tissue when exposed to AIV09, AIV48, and PR8. Infection with AIV07 was limited, with only the teat cistern of D1 and D2 showing infection. Only one donor (D2; Aberdeen Angus) was permissive to infection with H3N2, and infection to H1N1 was limited with only two donors showing NP-positive cells in the teat cistern and three donors in the gland cistern. D5, the only Holstein Friesian (dairy cow) in our study, was highly permissive to infection with both AIV09 and AIV48 infecting both the teat and gland cisterns. All donors from beef cattle breeds – Aberdeen Angus (D1, D2, and D6) and Limousin (D3 and D4) – were shown to be permissive to at least one virus in at least one tissue type. No consistent breed-restricted pattern of susceptibility was observed. Overall, these findings indicate that bovine mammary tissue supports infection by avian-adapted clade 2.3.4.4b H5N1 viruses, with susceptibility varying among donors and tissue sites.

**Table 3:** Permissiveness of cow teat and gland explants to infection with IAV. The table summarises the extent and pattern infection of each donor in this study to infection with each of the six viruses used and a mock for both the teat and gland cistern. Samples are reported as permissive (green box) if any infected cells were observed for all biological and technical replicates of that unique combination of donor, tissue, and virus. See Figure S3 for representative images of ‘focal’, ‘multifocal’, ‘multifocal/coalescing’, and ‘diffuse’ infection. Samples are only reported as non-permissive (grey box with ‘-’) if all biological and technical replicates of that unique combination of donor, tissue, and virus were negative for infection.

|  |  | Aberdeen Angus |  |  | Limousin |  | Holstein-Friesian |
| --- | --- | --- | --- | --- | --- | --- | --- |
|  |  | D1 | D2 | D6 | D3 | D4 | D5 |
| **AIV07** | Teat | Focal | Focal | – | – | – | – |
|  | Gland | – | – | – | – | – | – |
| **AIV09** | Teat | Multifocal | Multifocal | Multifocal | - | Multifocal | Multifocal |
|  | Gland | – | – | Focal | Focal | – | Multifocal |
| **AIV48** | Teat | Multifocal | Multifocal | – | Multifocal | – | Multifocal |
|  | Gland | – | – | Multifocal/coalescing | – | Focal | Focal |
| **H1N1** | Teat | – | - | – | – | Multifocal | Focal |
|  | Gland | – | Multifocal | Multifocal | – | Multifocal | – |
| **H3N2** | Teat | – | Multifocal | – | – | – | – |
|  | Gland | – | Focal | – | – | – | – |
| **PR8** | Teat | Multifocal | Multifocal/coalescing | Diffuse | Multifocal/coalescing | – | Multifocal |
|  | Gland | Diffuse | Multifocal | – | Multifocal | Multifocal/coalescing | Multifocal |
| **Mock** | Teat | – | – | – | – | – | – |
|  | Gland | – | – | – | – | – | – |

**Figure 2:**
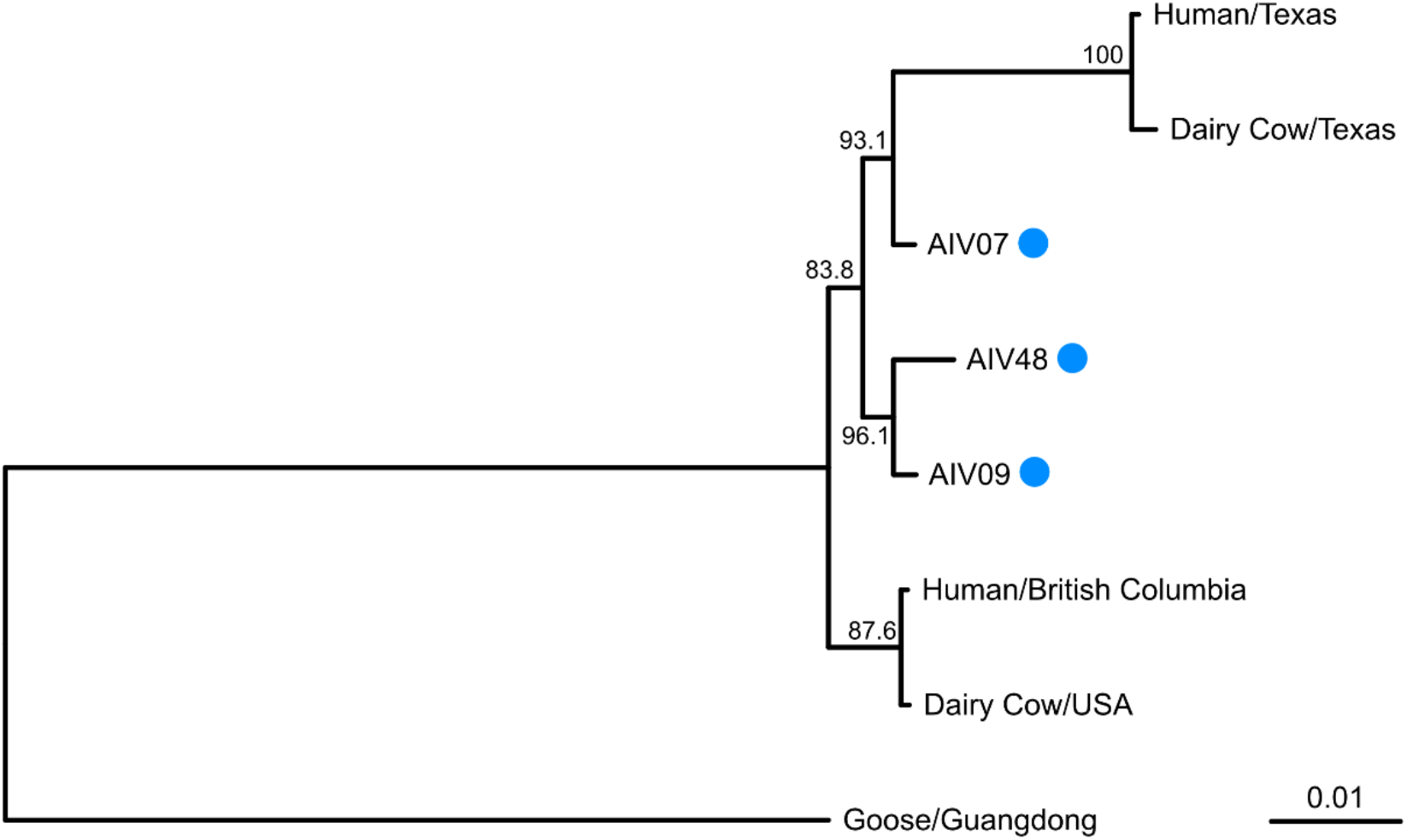
Phylogenetic tree of H5 genes used in this study. Maximum likelihood phylogeny of H5N1 HA sequences showing the evolutionary relationships between isolates used in this study and representative H5N1 viruses including bovine and human isolates from the ongoing H5N1 epizootic (A/dairy cow/Texas/24-008749-001-original/2024, A/Texas/37/2024, A/dairy cow/USA/002645-005/2025, and A/British_Columbia/PHL-2032/2024) and a prototypic pre-2020 H5N1 isolate (A/goose/Guangdong/1/1996). The tree was inferred using PhyML in Geneious under a GTR+G+I substitution model with four gamma rate categories. Bootstrap values are displayed on the nodes, the scale bar indicates substitutions per site, and blue circles denote viruses used in this study.

**Figure 3:**
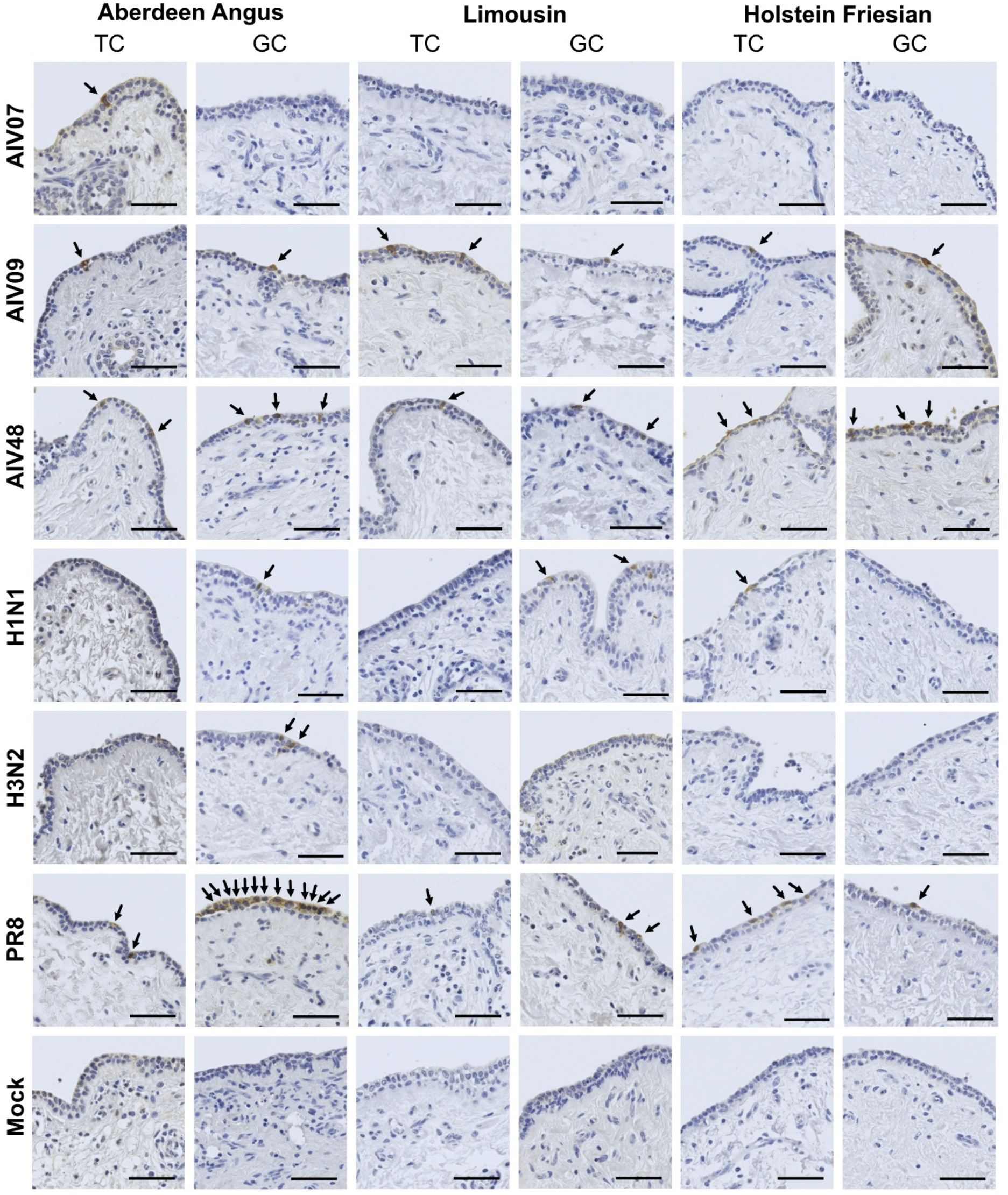
Immunohistochemistry staining results for one representative donor of each cattle breed. Each row represents tissue infected with one of six viruses or uninfected tissue (mock). Blue staining represents uninfected cell nuclei and infected cells appear as brown and are highlighted with black arrows.

## Discussion

The continued circulation of H5N1 in US dairy cattle, together with recent evidence of bovine exposure in the Netherlands, has raised concern about the risk of similar events in other countries with large cattle populations. This study aimed to assess the risk of avian-origin 2.3.4.4b H5N1 viruses infecting the mammary tissue of three common cattle breeds. Viral antigen was detected in explants from both beef (Aberdeen Angus, Limousin) and dairy (Holstein-Friesian) cattle, with no consistent breed-restricted pattern of susceptibility. Lectin staining showed abundant α2,6-linked and α2,3-linked sialic acids in epithelial cells across all donors and breeds. Together, these findings indicate that cattle breeds that are broadly used for food production can support infection by avian-origin H5N1 and may also be permissive to influenza A viruses with broader receptor usage.

The co-expression of α2,3- and α2,6-linked sialic acids in bovine mammary epithelium is notable because receptor usage is a major determinant of IAV host range. While avian viruses classically prefer α2,3-linked receptors and human-adapted viruses more often use α2,6-linked receptors, recent work suggests that bovine-associated H5N1 viruses may show at least limited or dual engagement of both receptor types [20,28,29]. Sustained viral replication within the bovine mammary gland may consequently represent a distinct multicellular niche in which selective pressures differ from those operating in respiratory tissues [23,29]. Even infrequent spillover events involving bovine mammary tissue may therefore provide conditions that permit viral adaptive change, allowing avian-adapted viruses to increase their binding to α2,6-linked sialic acid receptors and ability to infect humans. In parallel, the co-expression of α2,3- and α2,6-linked sialic acids in cow mammary glands may permit infection by viruses with broader receptor tropism and [30–32], where cattle are exposed to more than one influenza A virus, create opportunities for mixed infection and reassortment.

While large-scale outbreaks have thus far been reported primarily in dairy cattle in the US, our data indicate that the mammary tissue of common beef cattle is also permissive to infection. All cattle breeds used in this study are classed as international breeds by the Food and Agricultural Organization of the United Nations (FAO), meaning they are used widely outside the UK, including in countries such as USA, Australia, Germany, and Argentina [33]. Given the broad geographic circulation of H5N1 in wild birds, including along migratory flyways, susceptible cattle populations in multiple regions may be at risk of exposure. Consequently, differences in outbreak magnitude between countries may be more plausibly explained by differences in population structure and management practices than by intrinsic biological resistance of particular cattle breeds. More broadly, these findings support the possibility that influenza A virus infection in cattle is under-recognised. Historically, infection was regarded as very infrequent in cattle, but recent retrospective serology has identified nucleoprotein-reactive antibodies in 586 of 1,724 bovine sera across multiple breeds and physiological states, suggesting that prior exposure may be more common than previously appreciated [34].

Our findings show that mammary tissue from common cattle breeds is permissive to clade 2.3.4.4b H5N1 infection. These results support the view that bovine mammary epithelium represents a susceptible tissue environment and suggest that IAV infection in cattle may be more widespread than previously assumed. Accordingly, cattle should be incorporated into H5N1 surveillance frameworks, particularly during poultry outbreaks or incursions of infected wild birds. Surveillance strategies that combine PCR-based detection of active infection with serology for prior exposure may provide a more complete picture of virus circulation at both individual- and herd-level. Overall, integrating cattle into preparedness planning is a precautionary and evidence-based response to the expanding host range of clade 2.3.4.4b H5N1 viruses.

## Acknowledgements

The authors would like to thank the staff at Sandyford Abattoir (Paisley, Scotland) and the staff at the University of Glasgow Veterinary Diagnostic Services for their support with post-mortem services. The authors would specifically like to thank Colin Loney and Deirdre McLachlan from the Centre for Virus Research bioimaging team, in addition to Lynn Marion Stevenson, Lynn Oxford, Frazer Bell, and Lewis Kidd from the Veterinary Diagnostic Services Histology service for the support with FFPE tissue blocks, sectioning and mounting.

## Author contributions

Conceptualisation – RR, SKW, EH, PRM

Formal analysis - SKW

Funding acquisition – EH, PRM

Investigation – RR, SKW, HM, HC

Methodology – RR, SKW, HM

Supervision – EH, PRM

Visualisation – RR, SKW, HM

Writing (original draft) - RR, SKW

Writing (review and editing) - RR, SKW, EH, PRM

## Conflicts of interest

The authors declare no conflicts of interest.

## Funding information

R.R, S.K.W, H.M, E.H, and P.R.M were supported by the UK Medical Research Council (MRC) and Department for Environment, Food and Rural Affairs (DEFRA) through the consortium grant FluTrailMap-One Health [MR/Y03368X/1]. We acknowledge funding from MRC/UKRI [MR/Y03368X/1] to the CVR [MC_UU_0034/2 and MC_UU_0034/3] for P.R.M and [MC_UU_00034/1] for E.H. P.R.M also acknowledges funding by the Biotechnology and Biological Sciences Research Council [BB/V004697/1]. HC is funded by the University of Glasgow Vet Fund.

## Supplementary Data

**Table S1:**
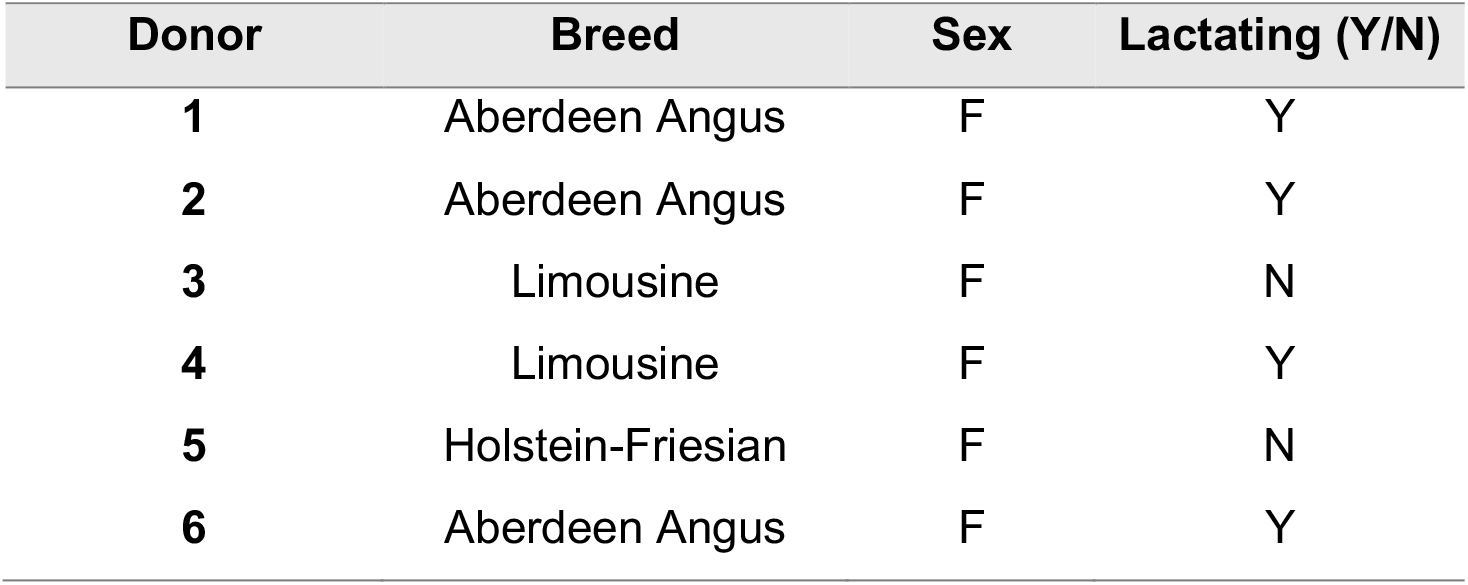
Table of breed, sex and lactation status of cattle.

| Donor | Breed | Sex | Lactating (Y/N) |
| --- | --- | --- | --- |
| 1 | Aberdeen Angus | F | Y |
| 2 | Aberdeen Angus | F | Y |
| 3 | Limousine | F | N |
| 4 | Limousine | F | Y |
| 5 | Holstein-Friesian | F | N |
| 6 | Aberdeen Angus | F | Y |

**Figure S1:**
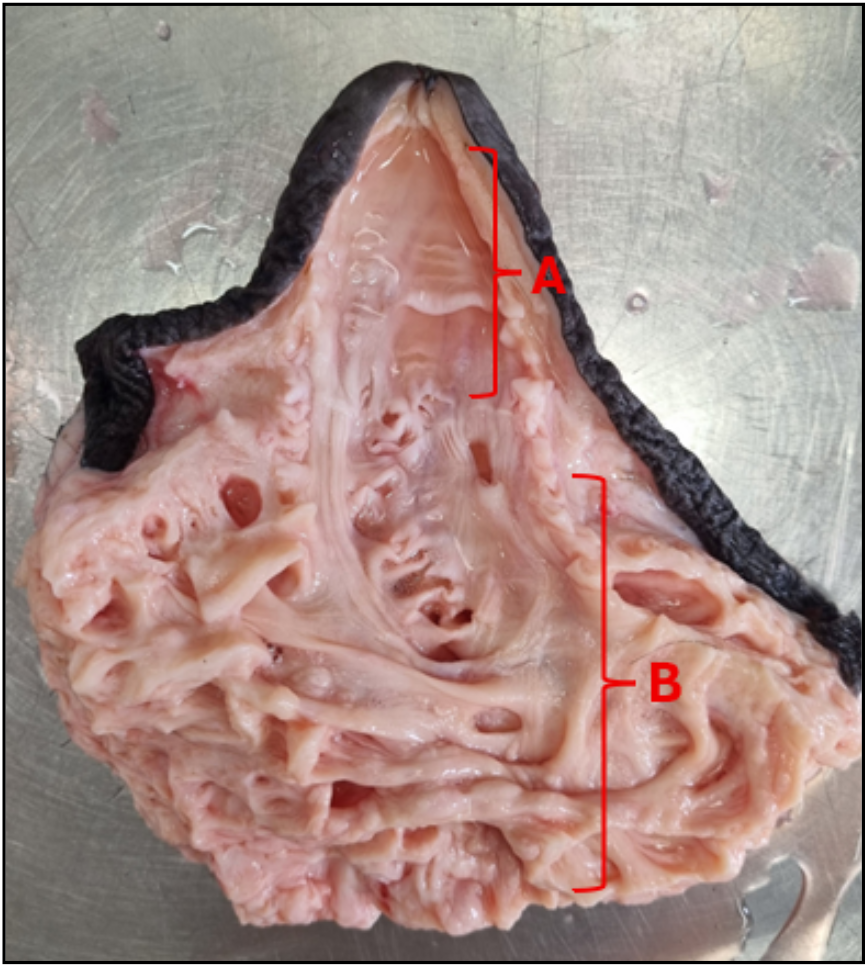
Sampling locations from the mammary glands. Cross section of teat cistern (A) and gland cistern (B) removed from mammary quarter.

**Figure S2:**
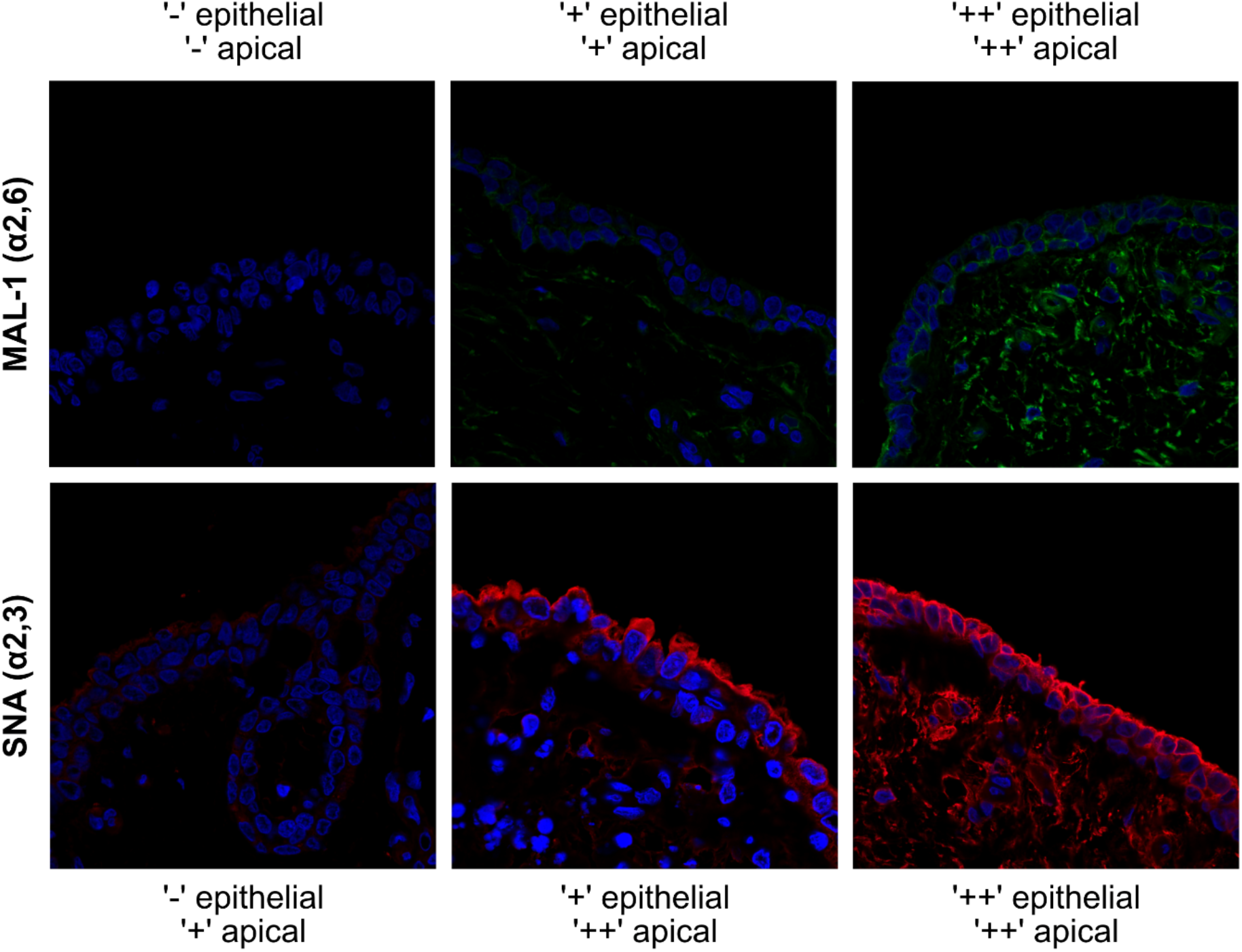
Representative images of each relative grade for the Lectin staining. Staining intensity was expressed by relative grades from negative (-) to weak (+) to strong (++). The above images show representative images of what each grade looked like. No captured samples were negative for SNA on the apical surface. MAL-1 = α2,3, SNA = α2,6.

**Figure S3:**
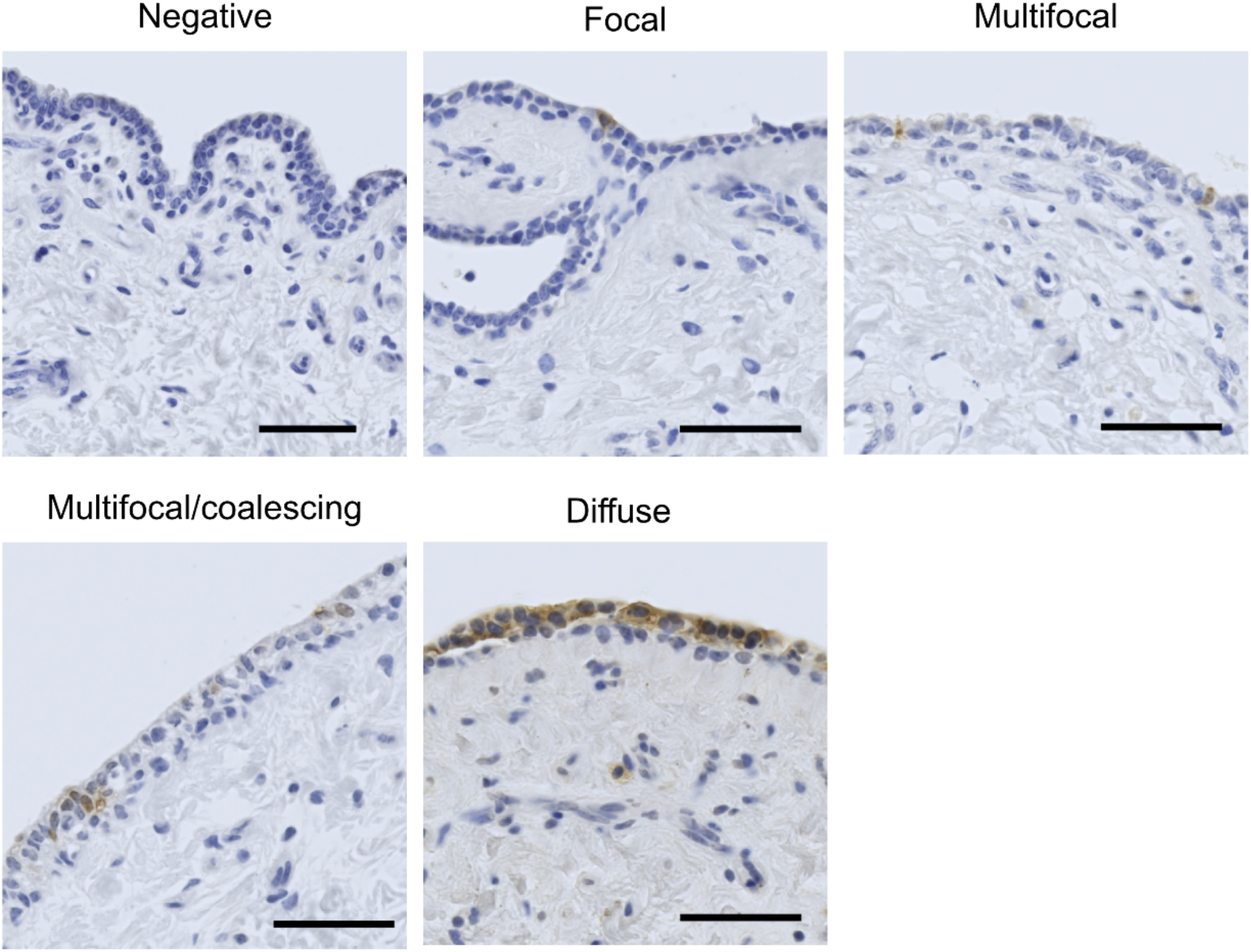
Types of infection reported in IHC staining. Representative images of each type of infection observed in the IHC staining: ‘negative’, ‘focal’, ‘multifocal’, ‘multifocal/coalescing’, and ‘diffuse’ infection.

## Notes

http://dx.doi.org/10.5525/gla.researchdata.2237

### Competing Interest Statement

The authors have declared no competing interest.

## References

1. Xu X, Subbarao K, Cox NJ, Guo Y. Genetic Characterization of the Pathogenic Influenza A/Goose/Guangdong/1/96 (H5N1) Virus: Similarity of Its Hemagglutinin Gene to Those of H5N1 Viruses from the 1997 Outbreaks in Hong Kong. Virology. 1999;261: 15–19. doi:10.1006/VIRO.1999.9820

2. Sims LD, Ellis TM, Liu KK, Dyrting K, Wong H, Peiris M, et al. Avian Influenza in Hong Kong 1997–2002. https://doi.org/101637/0005-2086-47.s3832. 2003;47: 832–838. doi:10.1637/0005-2086-47.S3.832

3. Lee DH, Bertran K, Kwon JH, Swayne DE. Evolution, global spread, and pathogenicity of highly pathogenic avian influenza H5Nx clade 2.3.4.4. J Vet Sci. 2017;18: 269–280. doi:10.4142/JVS.2017.18.S1.269

4. Peacock TP, Moncla L, Dudas G, VanInsberghe D, Sukhova K, Lloyd-Smith JO, et al. The global H5N1 influenza panzootic in mammals. Nature 2024 637:8045. 2024;637: 304–313. doi:10.1038/s41586-024-08054-z

5. Agüero M, Monne I, Azucena Sánchez O, Zecchin B, Fusaro A, Ruano MJ, et al. Highly pathogenic avian influenza A(H5N1) virus infection in farmed minks, Spain, October 2022. Eurosurveillance. 2023;28: 2300001. doi:10.2807/1560-7917.ES.2023.28.3.2300001;TIMESTAMP:1775735894367

6. Kareinen L, Tammiranta N, Kauppinen A, Zecchin B, Pastori A, Monne I, et al. Highly pathogenic avian influenza A(H5N1) virus infections on fur farms connected to mass mortalities of black-headed gulls, Finland, July to October 2023. Euro Surveill. 2024;29. doi:10.2807/1560-7917.ES.2024.29.25.2400063

7. Lindh E, Lounela H, Ikonen N, Kantala T, Savolainen-Kopra C, Kauppinen A, et al. Highly pathogenic avian influenza A(H5N1) virus infection on multiple fur farms in the South and Central Ostrobothnia regions of Finland, July 2023. Euro Surveill. 2023;28. doi:10.2807/1560-7917.ES.2023.28.31.2300400

8. Uhart MM, Vanstreels RET, Nelson MI, Olivera V, Campagna J, Zavattieri V, et al. Epidemiological data of an influenza A/H5N1 outbreak in elephant seals in Argentina indicates mammal-to-mammal transmission. Nature Communications 2024 15:1. 2024;15: 9516–. doi:10.1038/s41467-024-53766-5

9. Rimondi A, Vanstreels RET, Olivera V, Donini A, Lauriente MM, Uhart MM. Highly Pathogenic Avian Influenza A(H5N1) Viruses from Multispecies Outbreak, Argentina, August 2023. Emerg Infect Dis. 2024;30: 812–814. doi:10.3201/EID3004.231725

10. Tomás G, Marandino A, Panzera Y, Rodríguez S, Wallau GL, Dezordi FZ, et al. Highly pathogenic avian influenza H5N1 virus infections in pinnipeds and seabirds in Uruguay: Implications for bird-mammal transmission in South America. Virus Evol. 2024;10. doi:10.1093/VE/VEAE031

11. Gamarra-Toledo V, Plaza PI, Gutiérrez R, Inga-Diaz G, Saravia-Guevara P, Pereyra-Meza O, et al. Mass Mortality of Sea Lions Caused by Highly Pathogenic Avian Influenza A(H5N1) Virus. Emerg Infect Dis. 2023;29: 2553–2556. doi:10.3201/EID2912.230192

12. Leguia M, Garcia-Glaessner A, Muñoz-Saavedra B, Juarez D, Barrera P, Calvo-Mac C, et al. Highly pathogenic avian influenza A (H5N1) in marine mammals and seabirds in Peru. Nature Communications 2023 14:1. 2023;14: 5489–. doi:10.1038/s41467-023-41182-0

13. Caserta LC, Frye EA, Butt SL, Laverack M, Nooruzzaman M, Covaleda LM, et al. Spillover of highly pathogenic avian influenza H5N1 virus to dairy cattle. Nature 2024 634:8034. 2024;634: 669–676. doi:10.1038/s41586-024-07849-4

14. Pekar JE, Crespo-Bellido A, Lemey P, Bowman AS, Peacock TP, Ochoa JN, et al. Can H5N1 avian influenza in dairy cattle be contained in the US? Cell. 2026;189: 699–705. doi:10.1016/J.CELL.2025.12.033

15. House of Representatives of the States General. Parliamentary Document 28807, no. 323 | Overheid.nl > Official publications. 26 Jan 2026 [cited 9 Apr 2026]. Available: https://zoek.officielebekendmakingen.nl/kst-28807-323.html

16. Long JS, Mistry B, Haslam SM, Barclay WS. Host and viral determinants of influenza A virus species specificity. Nat Rev Microbiol. 2019;17: 67–81. doi:10.1038/S41579-018-0115-Z;TECHMETA

17. Burrough ER, Magstadt DR, Petersen B, Timmermans SJ, Gauger PC, Zhang J, et al. Highly Pathogenic Avian Influenza A(H5N1) Clade 2.3.4.4b Virus Infection in Domestic Dairy Cattle and Cats, United States, 2024. Emerg Infect Dis. 2024;30: 1335–1343. doi:10.3201/EID3007.240508

18. Baker AL, Arruda B, Palmer M V., Boggiatto P, Sarlo Davila K, Buckley A, et al. Dairy cows inoculated with highly pathogenic avian influenza virus H5N1. Nature 2024 637:8047. 2024;637: 913–920. doi:10.1038/s41586-024-08166-6

19. Halwe NJ, Cool K, Breithaupt A, Schön J, Trujillo JD, Nooruzzaman M, et al. H5N1 clade 2.3.4.4b dynamics in experimentally infected calves and cows. Nature 2024 637:8047. 2024;637: 903–912. doi:10.1038/s41586-024-08063-y

20. Song H, Hao T, Han P, Wang H, Zhang X, Li X, et al. Receptor binding, structure, and tissue tropism of cattle-infecting H5N1 avian influenza virus hemagglutinin. Cell. 2025;188: 919–929.e9. doi:10.1016/j.cell.2025.01.019

21. Cargnin Faccin F, Gay LC, Regmi D, Compton S, Mejías TD, Brondani JC, et al. Experimental infection and viral pathogenesis of a human isolate of H5N1 highly pathogenic avian influenza strain in Jersey cows. npj Veterinary Sciences 2026 1:1. 2026;1: 2–. doi:10.1038/s44433-025-00002-5

22. Shi J, Kong H, Cui P, Deng G, Zeng X, Jiang Y, et al. H5N1 virus invades the mammary glands of dairy cattle through ‘mouth-to-teat’ transmission. Natl Sci Rev. 2025;12. doi:10.1093/NSR/NWAF262

23. Pinto RM, Sharp CP, Beeson M, Pankaew N, Hassard JA, Moxom A, et al. The cow udder is a potential mixing vessel for influenza A viruses. bioRxiv. 2025; 2025.08.29.673079. doi:10.1101/2025.08.29.673079

24. Animal and Plant Health Agency. Detection of influenza A matrix gene by real-time Taqman® RT-PCR. 2020 Sep. Available: https://science.vla.gov.uk/fluglobalnet/Documents/english/protocol_Taqman.pdf

25. De Wit E, Spronken MIJ, Bestebroer TM, Rimmelzwaan GF, Osterhaus ADME, Fouchier RAM. Efficient generation and growth of influenza virus A/PR/8/34 from eight cDNA fragments. Virus Res. 2004;103: 155–161. doi:10.1016/j.virusres.2004.02.028

26. Turnbull ML, Zakaria MK, Upfold NS, Bakshi S, Magill C, Das UR, et al. The potential of H5N1 viruses to adapt to bovine cells varies throughout evolution. Nature Communications 2025 16:1. 2025;16: 11042–. doi:10.1038/s41467-025-67234-1

27. Magaki S, Hojat SA, Wei B, So A, Yong WH. An Introduction to the Performance of Immunohistochemistry. Methods Mol Biol. 2019;1897: 289–298. doi:10.1007/978-1-4939-8935-5_25

28. Eisfeld AJ, Biswas A, Guan L, Gu C, Maemura T, Trifkovic S, et al. Pathogenicity and transmissibility of bovine H5N1 influenza virus. Nature 2024 633:8029. 2024;633: 426–432. doi:10.1038/s41586-024-07766-6

29. Naveed A. The bovine mammary gland as a crucible for zoonotic influenza virus emergence: Receptor-mediated adaptation of HPAI H5N1 clade 2.3.4.4b. Archives of Virology 2026 171:3. 2026;171: 89–. doi:10.1007/S00705-026-06529-0

30. Kristensen C, Jensen HE, Trebbien R, Webby RJ, Larsen LE. Avian and Human Influenza A Virus Receptors in Bovine Mammary Gland - Volume 30, Number 9—September 2024 - Emerging Infectious Diseases journal - CDC. Emerg Infect Dis. 2024;30: 1907–1911. doi:10.3201/EID3009.240696

31. Ríos Carrasco M, Gröne A, van den Brand JMA, de Vries RP. The mammary glands of cows abundantly display receptors for circulating avian H5 viruses. J Virol. 2024;98. doi:10.1128/JVI.01052-24;WEBSITE:WEBSITE:ASMJ;WGROUP:STRING:PUBLICATION

32. Nelli RK, Harm TA, Siepker C, Groeltz-Thrush JM, Jones B, Twu NC, et al. Sialic Acid Receptor Specificity in Mammary Gland of Dairy Cattle Infected with Highly Pathogenic Avian Influenza A(H5N1) Virus - Volume 30, Number 7—July 2024 - Emerging Infectious Diseases journal - CDC. Emerg Infect Dis. 2024;30: 1361–1373. doi:10.3201/EID3007.240689

33. Food and Agriculture Organization of the United Nations. Domestic Animal Diversity Information System (DAD-IS. 2026 [cited 9 Apr 2026]. Available: https://www.fao.org/dad-is/browse-by-country-and-species/en/

34. Lang Y, Shi L, Roy S, Gupta D, Dai C, Khalid MA, et al. Detection of antibodies against influenza A viruses in cattle. J Virol. 2025;99: e02138–24. doi:10.1128/JVI.02138-24

